# Large scale comparative wheat phosphoproteome profiling pinpoints temperature-associated molecular signatures

**DOI:** 10.1101/2024.08.20.608123

**Authors:** Cássio Flávio Fonseca de Lima, Tingting Zhu, Lisa Van den Broeck, Brigitte Van De Cotte, Anna M. Locke, Rosangela Sozzani, Ive De Smet

**Author notes:** Rijk Zwaan, 4793 RS Fijnaart, The Netherlands. Protealis, B-9052 Gent, Belgium (Lisa Van den Broeck). Author for correspondence: Ive De Smet. The author responsible for distribution of materials integral to the findings presented in this article in accordance with the policy described in the Instructions for Authors (https://academic.oup.com/plcell/pages/General-Instructions) is: Ive De Smet.

## Abstract

Elevated temperatures resulting from climate change adversely affect natural and crop ecosystems, necessitating the development of heat-tolerant crops. We established a framework to precisely identify wheat protein-phosphorylated sites associated with varying temperature sensitivities. We identified specific kinases primarily associated with a specific temperature, but our results also suggest a striking overlap between cold and heat signaling. We furthermore demonstrated that the phosphorylation state of a specific set of proteins creates a unique signature for heat stress tolerance. These findings can potentially aid in the identification of targets for breeding or genome editing to enhance sub/supra optimal temperature tolerance.

## INTRODUCTION

Throughout their life cycle, plants encounter a range of temperatures, from freezing conditions to extreme heat. However, plants exhibit remarkable adaptability to these conditions (Anderson and Song 2020; Fernández-Marín et al. 2020; Praat et al. 2021; Ul Hassan et al. 2022). Nonetheless, the increasing trend of rising temperatures, driven by climate change, affects the productivity of natural and crop ecosystems (Wang et al. 2018; Doughty et al. 2023). An extreme increase in temperature above a critical threshold (heat stress) leads to irreversible damage to plant growth and development and can even result in plant death (Jagadish et al. 2021). Moreover, even a slight increase of only a few degrees above the optimal growth temperature can have a wide range of unfavorable effects at every developmental stage (Jacott and Boden 2020; Zhu et al. 2021).

In wheat, temperature sensitivity varies across different developmental stages and among different varieties. While the vegetative stage is generally more tolerant to temperature fluctuations than the reproductive stage, the specific thresholds for optimal and stress-inducing temperatures can differ depending on the variety (Porter and Gawith 1999). For instance, winter wheat is often sown in conditions where temperatures can drop as low as 8°C (Porter and Gawith 1999). Conversely, temperatures above 30°C are widely recognized as harmful, negatively affecting processes like photosynthesis (Porter and Gawith 1999) and reducing the activity of enzymes responsible for starch synthesis within the 30-40°C range (Keeling et al. 1994; Impa et al. 2020). Additionally, temperatures exceeding 34°C can inhibit chlorophyll biosynthesis, leading to accelerated leaf senescence (Asseng et al. 2013).

Given that air temperatures have been rising in most of the major cereal cultivating regions around the world (Zhao et al. 2017), breeding varieties that are tolerant to increased temperatures is a fundamental approach to deal with climate change and to assure future food security (Driedonks et al. 2016). To enable sustainable agriculture, additional breeding schemes are urgently needed. The current state-of-the-art plant breeding techniques mainly focus on genetic/genomic markers linked to desirable traits through, for example, genome-wide association studies (GWAS) (Spindel et al. 2016) or marker-assisted selection (MAS) (Hasan et al. 2021). However, these methods primarily focus on PCR-based capturing of DNA-level variations and do not directly capture post-translational modifications that regulate protein function, which are crucial for understanding complex traits like heat tolerance.

Phosphoproteomics provides a complementary approach by directly analyzing protein modifications that affect cellular functions and stress responses (Li and Yan 2020; Xu et al. 2023). Phosphorylation is a key post-translational modification that regulates protein activity, stability, and interactions. By identifying phosphorylation sites (phosphosites) associated with heat tolerance, we can uncover the dynamic regulatory networks that underpin this trait. Unlike approaches that target specific genetic sequences, proteomics offers a holistic view of protein regulation and activity in response to environmental changes.

In this study, we established a framework to precisely identify wheat protein-phosphorylated sites associated with varying temperature sensitivities. Our findings demonstrated that the phosphorylation state of a specific set of proteins creates a unique signature for heat stress tolerance. These findings will aid in the identification of targets for breeding or genome editing to enhance heat tolerance. In addition, we found a striking overlap between cold and heat signaling, that will be the basis for future functional characterization of common signaling components.

## RESULTS

### Phenotypic plasticity in wheat at different temperatures

Vegetative development displays high plasticity, namely phenotypic modifications to maintain stable equilibrium or fitness (Schneider 2022). To explore the phenotypic plasticity of the vegetative organs in wheat, we focused on the juvenile phase. We analyzed growth (length of 1^st^, 2^nd,^ and 3^rd^ leaves) and development (reaching the 13^th^ stage on the Zadoks scale) along a temperature gradient (14, 19, 24, 29 and 34 °C) in 14-day-old seedlings from eight wheat varieties with distinct sensitivities to high temperatures (**Fig. 1A-B, Extended data Fig. 1-2 and Extended data tables 1-2).** For the vegetative stage of wheat, the optimal temperature for growth is within the range of 20-25°C (Zhu et al. 2021). Therefore, we used 14°C and 34°C as temperatures that induce chilling and heat stress, respectively, while 19°C and 29°C are just outside that optimum. Our results suggested that the three leaves responded differently to the temperature gradient and exhibited organ-specific plasticity (**Fig. 1C, Extended data Fig. 3).** Based on the available data, we estimated that the maximum length of the second leaf occurs around 21.6°C, while the first and third leaves reach their greatest lengths at approximately 17.7°C and 27.4°C, respectively (**Extended data Fig. 3a, b**). Furthermore, from the 1^st^ to the 3^rd^ leaf, the growth response along the temperature gradient became increasingly uniform for all varieties (**Fig. 1C**). The transition from stage Z12 to Z13 was strongly repressed at 14 °C and peaked between 19 °C and 29 °C, and the high temperature-sensitive varieties showed a very sharp decrease at 34 °C (**Fig. 1D)**. Plasticity at the level of organ length and developmental stage was associated with previously determined temperature-sensitive or (moderately) tolerant responses (**Fig. 1E, Extended data table 1)**. In summary, there is an organ-specific threshold at which supra-optimal temperatures are no longer growth/development-promoting, but instead repress growth/development.

**Figure 1.**
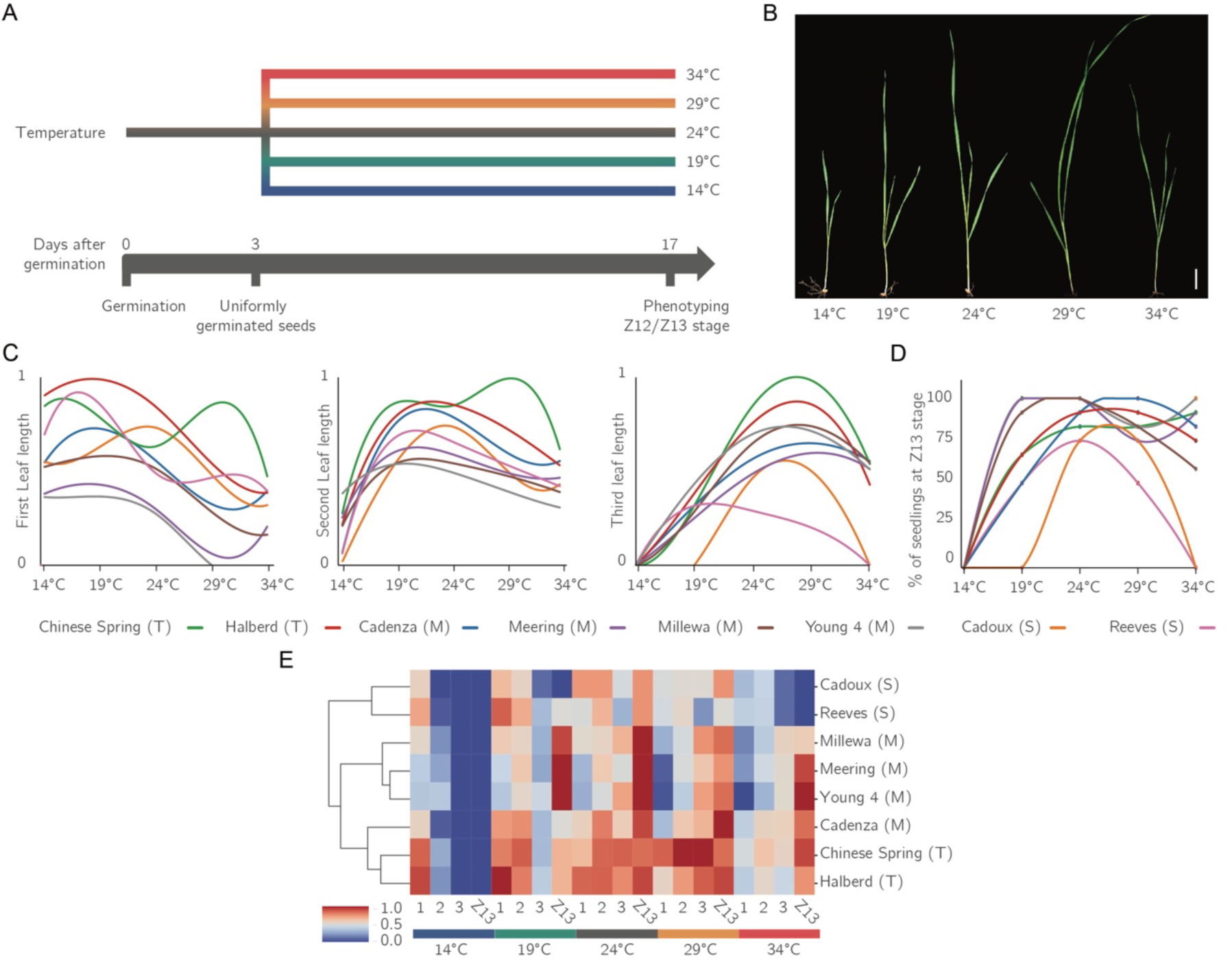
Growth and developmental responses of wheat (*Triticum aestivum* L.) along the temperature gradient. **A.** Phenotyping setup. Developmental stage and leaf (first, second, or third) lengths were measured for 14 days after germination (DAG) of seedlings continuously grown under five distinct temperature conditions (14, 19, 24, 29, and 34 °C). **B.** Representative pictures of 14 DAG wheat var. Cadenza treatment at the indicated temperatures. **C**. Graphs of the length of the first (1), second (2) or third leaf (3) of the indicated wheat varieties after exposure to the temperature gradient, values were scaled to the shortest/longest (0-1) measurement of each leaf per variety. Third-degree polynomial curves were fitted to the normalized length data. Raw values were normalized for 0-1 min/max. Sensitive (S), moderately tolerant (M) or tolerant (T) to temperature stress is indicated. **D**. Graph depicting the percentage of seedlings at Z13 development stage (Zadoks scale) of the indicated wheat varieties after exposure to the temperature gradient. **E**. Heatmap compiling normalized measured leaf responses adjusted per first (1), second (2) or third leaf (3) or percentage of seedlings that reached Z13 scale, color range adjusted to 0-1 for both measurement values (from 1c) and percent seedlings at Z13 (from 1d).

### Phosphoproteomes of wheat variety Cadenza at different temperatures

Because distinct varieties respond differently, we hypothesized that these threshold responses are molecularly encoded and fine-tuned differently in these varieties. Protein (de)phosphorylation pathways are dynamic, complex, and interconnected, and play a crucial role in signal transduction to regulate multiple biological processes (Derouiche et al. 2012). To identify molecular signatures associated with plasticity and the critical temperature threshold, we first examined protein phosphorylation dynamics in the second leaf of two-week-old wheat seedlings exposed to each temperature along the same gradient for 60 minutes (**Fig. 2A**). For this, we used the variety “Cadenza”, because the available genetic resources (Krasileva et al. 2017) make this variety well-suited for future functional follow-up studies.

**Figure 2.**
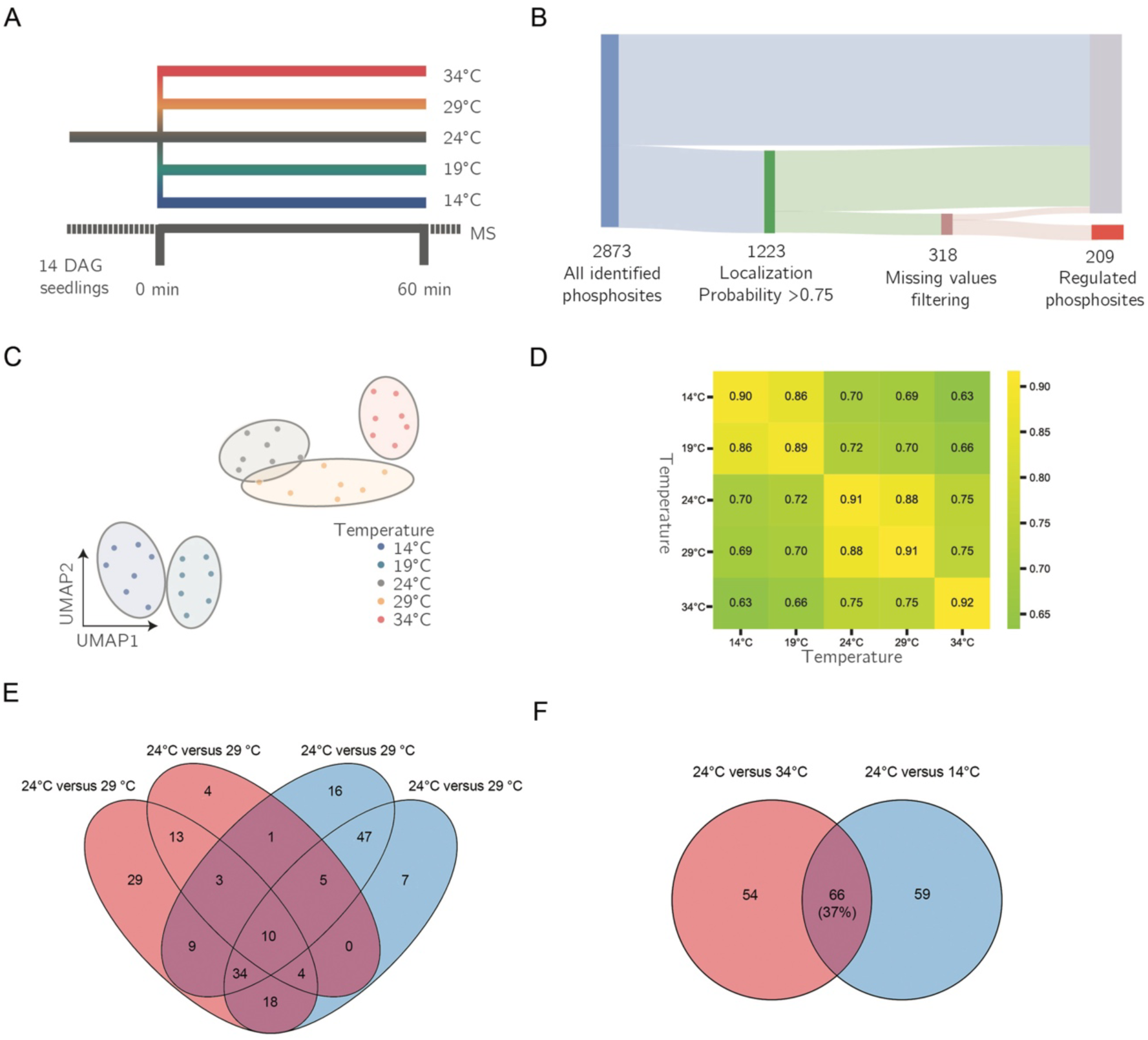
Molecular responses of wheat (*Triticum aestivum* L.) to a temperature gradient. **A**. Phosphoproteomic workflow. Cadenza seedlings were continuously grown at the control temperature (24 °C) until 14 DAG and were transferred to one of the indicated temperatures for 60 min. The second leaf was harvested and subjected to our LC-MS/MS workflow. **B**. Phosphoproteomics results for Cadenza along the temperature gradient, indicating the phosphorylated sites and filtering steps. **C**. Uniform manifold approximation and projection (UMAP) plot (Euclidean distance, n=3) of the phosphoproteome replicates using all identified phosphorylated sites, indicating the five temperature groups. **D**. Euclidean distance matrices grouped by biological replicates at distinct temperatures. **E-F.** Overlap of significant phosphorylated sites between all different temperatures (E) and between the two extreme temperatures (F).

Out of the 2873 identified phosphosites, we retained 1223 high-quality phosphosites for further analyses (**Fig. 2B** **and Extended data tables 2-3**). The highly reproducible phosphosite intensity profiles separated into three temperature-specific groups: a sub-optimal temperature group (14 °C and 19 °C), an optimal or warm temperature group (24 °C and 29 °C), and a heat-stress group (34 °C) (**Fig. 2C-D**). Subsequent analyses focused on 209 phosphosites that were significantly affected by temperature (**Fig. 2B**). When comparing the significant phosphosites at each temperature with respect to the 24 °C control, we observed a striking overlap between the lower (14 °C and 19 °C) and higher temperatures (29 °C and 34 °C) (**Fig. 2E-F**). This is a strong indication that high and low temperature signaling components overlap. We inferred two clusters that followed an increasing (cluster “up”) or decreasing phosphorylation trend (cluster “down”) along the temperature gradient and those two clusters displayed unique gene ontology signatures (**Extended data Fig. 4a-c and Extended data table 3**). Next, we sub-classified the two main clusters into 12 sub-clusters (**Fig. 3A-B)** and we grouped these according to their phosphoprofiles, namely a “bell-shaped”, “late” or “gradual” response. To identify phosphorylated proteins associated with plasticity and/or temperature threshold, we focused on (inverted) bell-shaped profiles along the temperature gradient (clusters 4, 7, 8, and 12) and late responses (clusters 1, 2, 3, and 9) (**Fig. 3B**). These showed a unique enrichment in nucleosomal DNA binding and photosynthesis-associated processes, respectively (**Fig. 3C**). In conclusion, a brief temperature treatment induced distinct phospho-signatures that revealed potential regulators of plasticity and/or temperature threshold.

**Figure 3.**
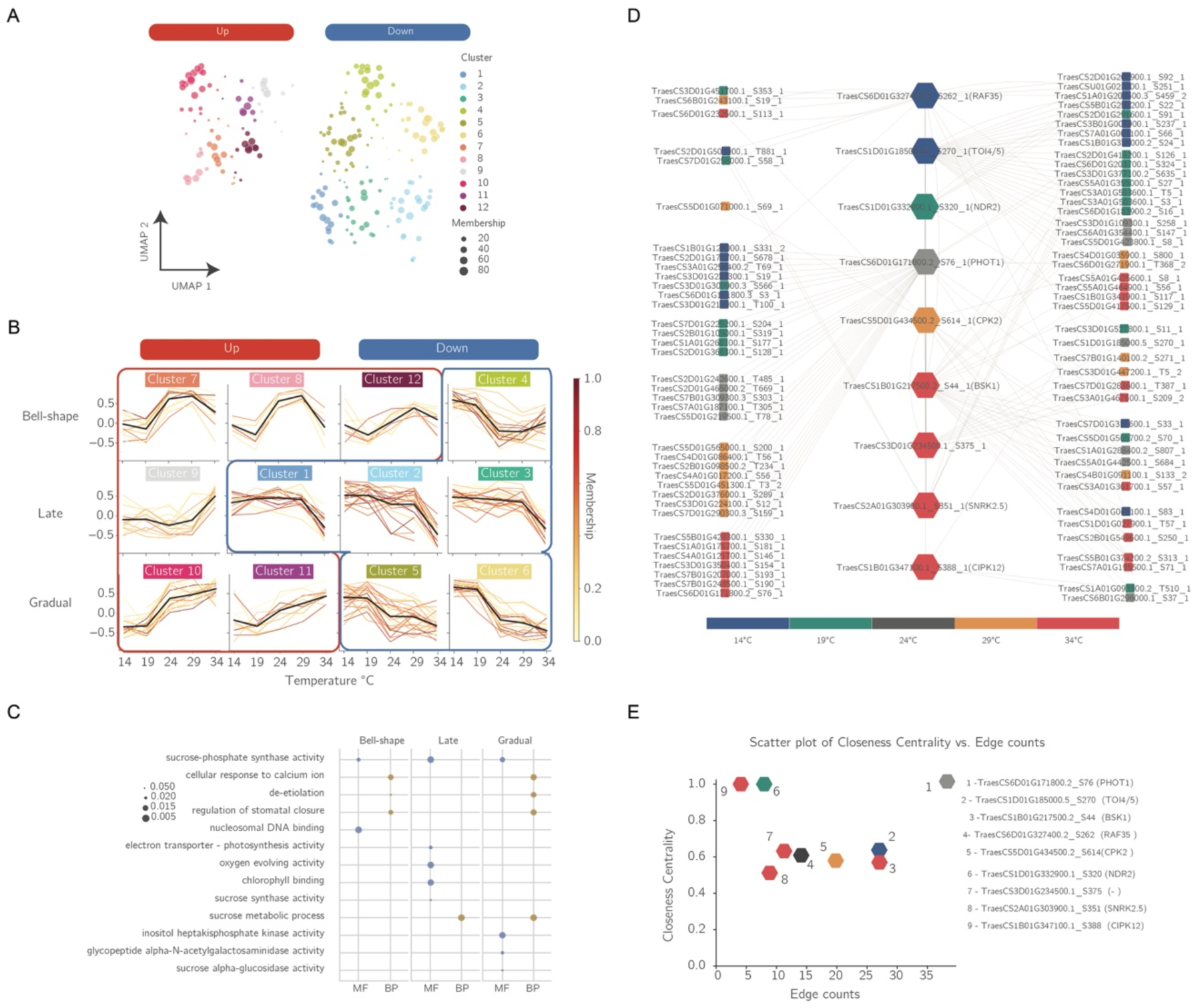
Signaling network of wheat at different temperatures. **A.** UMAP clusters of 209 temperature-responsive phosphorylated sites. Two main clusters (“Up” and “Down”) are indicated and those are further subdivided into 12 subgroups. Dot sizes represent the membership of a particular phosphorylated site in the respective cluster map. **B**. Bell-shaped, late and gradual temperature response profiles. **C.** Gene Ontologies associated with three indicated groups. **D.** Dynamic Bayesian networks of temperature gradient phosphoproteomes, showing nine kinases (hexagons, with the name of the best-hit Arabidopsis orthologue between parentheses) predicted to be tightly linked with indicated downstream proteins (circles). Colors indicate the peak of phosphorylation for the labelled site along the gradient of temperature. **E.** Relative importance of kinases in the predicted dynamic Bayesian network given their closeness centrality (CC) value and number of nodes (ND) connected (CC*ND).

**Figure 4.**
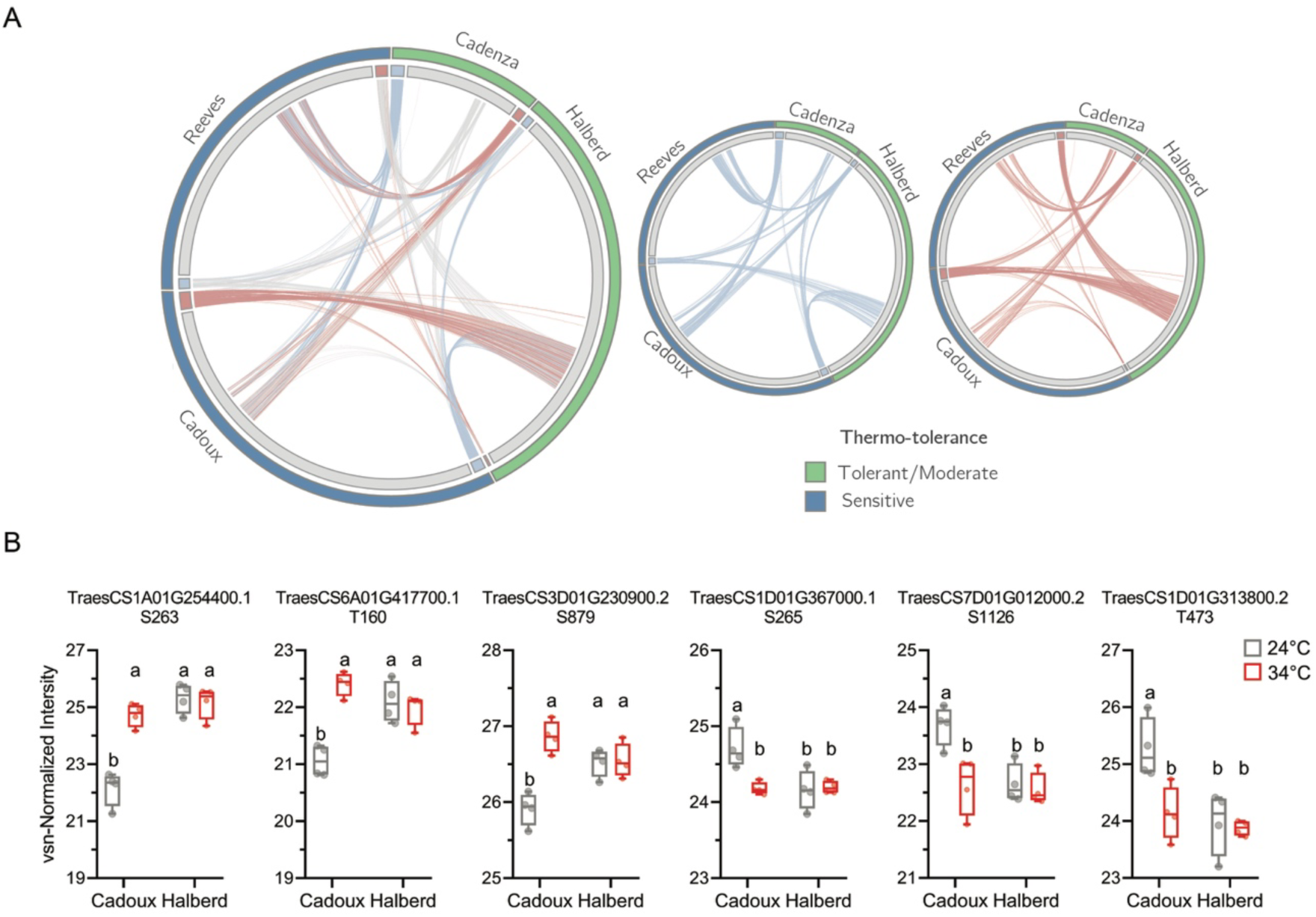
Molecular patterns of wheat varieties grown at different temperatures. **A.** Circos plot visualizing the regulation of phosphorylated sites in two temperatures (24-34 °C) in four wheat varieties (Cadenza, Halberd, Reeves and Cadoux). Lines represent a single phosphorylated site connects identifications of distinct varieties. Colors represent regulation in a certain variety and their status in another (blue, down-regulated; red, upregulated and grey, not-significantly affected). **B.** Representative examples of protein phosphorylation status and change at control (24 °C) and elevated temperature (34 °C) in Cadoux (sensitive) and Halberd (tolerant) plants. Box plots with individual data points represent the distribution of individual replicates, showing the median with Tukey-based whiskers. Letters indicate significant differences based on two-way ANOVA and post-hoc multiple comparisons tests (*p* < 0.05) between genotypes and between 24 °C and 34 °C.

### Specific kinases regulate signaling at distinct temperatures

To identify kinases that control plasticity and the critical temperature threshold, we employed a dynamic Bayesian network approach (Madison et al. 2023) using the high-confidence and consistently quantified phosphosites from the Cadenza temperature gradient phosphoproteome. We predicted nine kinases associated with the dynamic phosphorylation profiles of 104 phosphorylated sites within 95 proteins (**Fig. 3D, Extended data Fig. 5, Extended data table 4)**. To rank these kinases, we examined their edge counts and closeness centrality in the network (**Fig. 3E**). This pinpointed PHOTOTROPIN 1 (PHOT1) as an important regulatory kinase along the temperature gradient (**Fig. 3D-E**). In Arabidopsis, PHOT1 regulates photosynthetic activity by regulating phototropism, chloroplast relocation and stomatal opening (Wang et al. 2021). Moreover, a wheat ortholog of the Arabidopsis MITOGEN ACTIVATED PROTEIN KINASE KINASE KINASE 6 (MAP4K6)/TOT3-INTERACTING PROTEIN 4 (TOI4) and MAP4K5/TOI5 is the second most connected protein (**Fig. 3E**). Furthermore, wheat TOI4/5 had one of the highest scores for feature importance along the temperature gradient, suggesting a key role in temperature signaling (**Extended data table 5**). Indeed, in Arabidopsis and wheat, TARGET OF TEMPERATURE (TOT3), TOI4 and TOI5 are involved in thermomorphogenesis (Vu et al. 2021) and in soybean, TOI5 and TOT3 were suggested to play a central role in cold signaling (Van den Broeck et al. 2023).

### A phosphoprotein-based signature for heat tolerance

Similar to protein abundance (Vítámvás et al. 2019; Mahmood et al. 2022), we hypothesized that protein phosphorylation status could be used as a biomarker for determining (a)biotic stress tolerance or sensitivity in crops. To assess whether different wheat varieties (specifically thermosensitive versus thermotolerant ones) display a common or distinct protein phosphorylation status and change, we analyzed the highly reproducible phosphoproteomes at 24 °C and 34 °C of – in addition to Cadenza (39 up / 83 down regulated sites at 24 °C versus 34 °C) – three additional varieties with previously reported distinct high-temperature sensitivities (**Fig. 1C-E, Extended data Fig. 1**), namely Cadoux (24 up / 33 down), Halberd (6 up / 34 down) and Reeves (24 up / 23 down) (**Extended data Fig. 6a-b, Extended Data table 6**). Four phosphosites displayed up/downregulation in both sensitive varieties, but these were not regulated in the (medium) tolerant varieties (**Fig. 3A**). In addition, 59 phosphosites displayed up/down regulation in one of the sensitive varieties, but these were not regulated in the (medium) tolerant varieties (**Fig. 3A**). These differentially phosphorylated sites support the idea that specific protein phosphorylation status and response are associated with the sensitive and tolerant nature of a variety. We compared Cadoux and Halberd to pinpoint phosphosites that could explain high temperature tolerance. To reveal the protein phosphorylation status potentially associated with heat tolerance, we identified phosphosites that were up-or downregulated in one variety, not regulated in the other variety, and already at a high/low level in the control condition (**Fig. 3B and Extended data table 6-7**). In conclusion, leveraging the phosphoproteomes of tolerant and sensitive varieties allowed the identification of putative phosphoprotein-based biomarkers that can be used as predictive and diagnostic markers in wheat (**Fig. 3B**).

## DISCUSSION

We previously proposed the concept that responsiveness to temperature should be viewed from the context of dose dependency instead of a binary process (Zhu et al. 2022). Our growth, development, and molecular data provide evidence to support this concept, revealing key regulatory kinases and temperature-dependent processes. In addition, identifying post-translationally regulated players in the temperature response is crucial for functional studies and the identification of breeding markers or genome editing targets.

By comparing phosphoproteomes at low and high temperatures, we revealed a striking overlap between the signaling cascades associated with both temperature extremes. This provides the basis to explore such common regulators in more detail. In addition, we identified several novel phosphoproteins associated with heat tolerance in wheat. These phosphoproteins play crucial roles in various cellular processes that contribute to the plant’s ability to withstand high temperatures. Understanding the specific functions of these phosphoproteins can provide insights into their mechanisms of action and their potential as targets for crop improvement. One key group of phosphoproteins identified is involved in photosynthesis and energy metabolism. For example, the phosphorylation state of proteins related to the light reactions of photosynthesis, such as those in the thylakoid membrane, can influence the efficiency of photosystem II and protect the plant from photodamage under heat stress. By maintaining efficient photosynthetic activity, these proteins help ensure that the plant continues to produce the energy needed for growth and survival during high-temperature conditions. Another significant group includes phosphoproteins involved in signal transduction pathways, particularly those related to heat shock response and protein homeostasis. These proteins can modulate the activity of heat shock factors and other transcriptional regulators that activate the expression of heat shock proteins (HSPs). HSPs function as molecular chaperones, stabilizing proteins and preventing aggregation under stress conditions, thereby enhancing the plant’s resilience to heat. Additionally, phosphoproteins involved in cell wall remodeling and membrane stability were also identified. The phosphorylation of these proteins may regulate the structural integrity of cell walls and membranes, helping to maintain cellular function and prevent damage caused by heat-induced dehydration and oxidative stress.

In this study, we demonstrate a proof-of-concept for an experimental framework to identify key phosphosites by comparing crop varieties with contrasting stress responses. By elucidating the roles of these novel phosphoproteins, we can better understand the complex regulatory networks that underpin heat tolerance in wheat. Ultimately, the phosphorylation state of selected proteins defines a phosphoprotein signature, and these protein phosphorylation states serve as biomarkers for heat stress tolerance. This knowledge can inform breeding strategies and the development of biotechnological approaches aimed at enhancing heat tolerance in crops, ultimately contributing to food security in the face of climate change.

## METHODS

### Wheat plant materials and temperature treatment/growth conditions

We used various bread wheat varieties (*T. aestivum*, AABBDD), including those with previously reported high-temperature sensitivities (Mason et al. 2010; Maphosa et al. 2014; Shirdelmoghanloo 2015; Browne et al. 2021) (**Extended Data table 1**). Seeds were embedded in wet paper, enclosed in plastic wrap, and stratified at 4 °C for three continuous days. Subsequently, the seeds were transferred to room temperature for an additional two days to initiate germination. Germinated wheat seeds with a radicle length of approximately 3 mm were manually selected and individually planted in a soil mixture (50% white peat, 40% garden peat, and 10% French coniferous bark enriched with biofertilizer - NPK ratio of 6:5:7, at a concentration of 8.0 kg/m³, with pH range between 5.0 and 6.5). Seedlings (one per pot) were cultivated in round plastic pots (6 × 6 cm) in an incubator set at one of the following temperatures: 14, 19, 24, 29, or 34 °C. The incubator maintained a 16-hour light/8-hour dark cycle, providing (100 μE m^-2^s^-1^ photosynthetically active radiation) from cool-white fluorescent tungsten tubes (Osram). Air humidity was maintained between 65% and 75% throughout the 14-day growth period. For the growth and development experiments, after the 14-day treatment, the seedlings were transferred to a new tray approximately 3 h after the start of the light period (∼9 am). Samples were carefully removed from the soil for photography. Each seedling was photographed using a Nikon D5300 camera with a ruler. The images were subsequently processed using ImageJ v1.53 (Schneider et al. 2012) for organ measurement, and the seedlings were classified according to the Zadok scale for cereal developmental (ZADOKS et al. 1974). In the current study, all seedlings fell within the 12^th^ (fully emerged second leaf) and 13^th^ stages (fully emerged third leaf). Twelve separate biological replicates were collected for each variety and temperature treatment. For phosphoproteomics experiments, the treatment of seeds and seedlings followed the same initial steps as in the growth and development experiments. However, in this study, Cadenza variety seedlings were grown only at 24 °C (control temperature) for the entire 14-day period. On the final day, the seedlings were exposed to short-term temperature treatment. Pots were randomly selected and transferred to incubators set to 14, 19, 24, 29, _-2 -1_ or 34 °C under constant light conditions (100 μE m s photosynthetically active radiation) for 60 min.

### Cubic spline interpolation

To estimate the optimal temperature for leaf elongation in our study, we used cubic spline interpolation to model the growth response of each leaf (first, second, and third) to the temperature gradient. By smoothing the observed data points, this method allowed us to interpolate and identify the temperature at which each leaf reached its maximum length. These estimated temperatures are based on the available data and serve as informed approximations, acknowledging the limitations inherent in extrapolating from a finite dataset.

### Protein extraction and phosphopeptide enrichment

Following the temperature treatment, the second leaf of each seedling was harvested using scissors and pooled with leaves from two other seedlings into a single 1.5 mL tube, forming a pooled sample. To account for the high biological variability and ensure technical precision we utilized a comprehensive approach. For each temperature condition (14 °C, 19 °C, 24 °C, 29 °C, and 34 °C), we collected samples from seven biological replicates of wheat seedlings, where each biological replicate was a pool of three individual seedlings. This pooling approach resulted in a total of seven biological samples per temperature condition. This design ensures that our dataset captures biological variability. Each sample pool was immediately frozen in liquid nitrogen and used as a biological replicate. For the varieties Halberd, Cadoux, and Reeves, similar procedures were repeated, but with only 21 °C and 34 °C treatments and four biological replicates per variety-treatment combination. After collection, all samples were stored in a -70 °C freezer for subsequent processing.

Total protein was extracted from all leaf samples according to our previously described procedure with minor modifications (Vu et al. 2017). One gram of finely ground leaf material was suspended in homogenization buffer containing 30% sucrose, 50 mM Tris-HCl buffer (pH 8), 0.1 M KCl, 5 mM EDTA, and 500 mM DTT in Milli-Q water, to which the appropriate amounts of the cOmpleteTM protease inhibitor mixture (Roche) and the PhosSTOP phosphatase inhibitor mixture (Roche) were added. The samples were sonicated on ice and centrifuged at 4 °C for 15 min at 3220 g to remove debris. The supernatants were collected, and methanol/chloroform precipitation was carried out by adding 3, 1, and 4 volumes of methanol, chloroform, and water, respectively. The samples were centrifuged at room temperature for 10 min at 3220 × g, and the aqueous phase was removed. After adding four volumes of methanol, the proteins were pelleted by centrifugation for 10 min at 3220 g. The pellets were washed with 80% acetone and centrifuged for 10 min at room temperature. The supernatants were discarded, and the pellets were air-dried. Protein pellets were resuspended in 8 M urea in 50 mM triethylammonium bicarbonate (TEAB) buffer (pH 8). Alkylation of cysteines was carried out by adding tris(carboxyethyl)phosphine (TCEP, Pierce) and iodoacetamide (Sigma-Aldrich) to final concentrations of 15 mM and 30 mM, respectively, and the samples were incubated for 15 min at 30 °C in the dark. Three mg of protein material was pre-digested with MS-grade lysyl endopeptidase (Wako Chemicals) for 4 h at 37 °C at an enzyme-to-substrate ratio of 1:300 (w:w). The mixtures were diluted 8-fold with 50 mM TEAB, followed by overnight digestion with trypsin (Promega) at an enzyme-to-substrate ratio of 1:100. The digest was acidified to pH 3 with trifluoroacetic acid (TFA) and desalted using SampliQ C18 SPE cartridges (Agilent), according to the manufacturer’s guidelines. For phosphopeptide enrichment, the desalted peptides were fully dried in a vacuum centrifuge and then resuspended in 500 μl of loading solvent [80% (v/v) acetonitrile, 6% (v/v) TFA]. Phosphopeptides were enriched as previously described (Vu et al. 2017). The resuspended peptides were incubated with 1 mg MagReSyn® Ti-IMAC microspheres for 20 min at room temperature with continuous mixing. The microspheres were washed once with wash solvent 1 (60% acetonitrile, 1% TFA and 200 mM NaCl) and twice with wash solvent 2 (60% acetonitrile and 1% TFA). The bound phosphopeptides were eluted with three volumes (80 μl) of elution buffer (40% acetonitrile and 1% NH_4_OH), immediately followed by acidification to pH 3 using 100% formic acid. Prior to MS analysis, the samples were vacuum-dried and re-dissolved in 50 μl of 2% (v/v) acetonitrile and 0.1% (v/v) TFA.

### LC-MS/MS analysis

The peptides were re-dissolved in 50 µL loading solvent A (0.1% TFA in water/ACN (98:2, v/v)), of which 3 µL was injected for LC-MS/MS analysis on an Ultimate 3000 RSLC nano LC (Thermo Fisher Scientific, Bremen, Germany) connected to a Q Exactive mass spectrometer (Thermo Fisher Scientific). The peptides were first loaded onto a µPAC™ Trapping column with C18-endcapped functionality (Pharmafluidics, Belgium), and after flushing from the trapping column the peptides were separated on a 50 cm µPAC™ column with C18-endcapped functionality (Pharmafluidics, Belgium) maintained at a constant temperature of 35 °C. Peptides were eluted by a linear gradient from 98% solvent A’ (0.1% formic acid in water) to 55% solvent B′ (0.1% formic acid in water/acetonitrile, 20/80 (v/v)) for 120 min at a flow rate of 300 nL/min, followed by a 5 min wash reaching 99% solvent B’. The mass spectrometer was operated in data-dependent positive ionization mode, automatically switching between MS and MS/MS acquisition for the five most abundant peaks in a given MS spectrum. The source voltage was set to 3 kV and the capillary temperature was 275 °C. One MS1 scan (m/z 400−2,000, AGC target 3 × 106 ions, maximum ion injection time 80 ms), acquired at a resolution of 70,000 (at 200 m/z), was followed by up to five tandem MS scans (resolution 17,500 at 200 m/z) of the most intense ions fulfilling predefined selection criteria (AGC target 5 × 104 ions, maximum ion injection time 80 ms, isolation window 2 Da, fixed first mass 140 m/z, spectrum data type: centroid, under-fill ratio 2%, intensity threshold 1.3xE4, exclusion of unassigned, 1, 5-8, >8 positively charged precursors, peptide match preferred, excluding isotopes on, dynamic exclusion time 12 s). The HCD collision energy was set to 25% of the normalized collision energy, and the polydimethylcyclosiloxane background ion at 445.120025 Da was used for internal calibration (lock mass).

### Database searching and quality control

MS/MS spectra were searched against the IWGSC RefSeq v1.0 database for *Triticum aestivum var.* Chinese spring (International Wheat Genome Sequencing Consortium (IWGSC) 2014) with MaxQuant software (version 1.6.10.43), a program package allowing MS1-based label-free quantification acquired from Orbitrap instruments (Cox and Mann 2008; Cox et al. 2014). The precursor mass tolerance was set to 20 ppm for the first search (used for nonlinear mass re-calibration) and to 4.5 ppm for the main search. Trypsin was used as the enzyme setting. Cleavages between lysine/arginine-proline residues and up to two missed cleavages were allowed. S-Carbamidomethylation of cysteine residues was selected as a fixed modification, and oxidation of methionine residues was selected as a variable modification. The false discovery rate for peptide and protein identification was set to 1% and the minimum peptide length was set to 7. The minimum score threshold for both the modified and unmodified peptides was set to 30. The MaxLFQ algorithm allowing for label-free quantification (Cox et al. 2014) and the “matching between runs” feature were enabled. To calculate the protein ratios, both unique and razor peptides (non-unique peptides assigned to a protein group with the largest number of identified peptides) were selected.

To process and analyze the dataset, we used a custom Python script executed within a Jupyter notebook to enable a reproducible research workflow (Kluyver 2016). The Jupyter notebook containing all codes and outputs used in this study is available on https://github.com/Cassio-Lima/Fonseca_de_Lima_2023. The ‘Phospho (STY) Sites.txt’ tabular data file was loaded into the Jupyter notebook for analysis. The specific Python libraries used for data loading and processing included Pandas (McKinney 2010), Numpy (Harris et al. 2020), PaDuA (Ressa et al. 2019), and Scipy (Virtanen et al. 2020) and visualization on Seaborn (Waskom 2021). Phosphorylated sites identified as reverse, potential contaminants, or with a localization probability lower than 75% were removed for subsequent analysis. Next, quantifiable multiplicity sites were expanded to three, and the intensity values obtained were transformed using variance normalizing stabilization (vsn) (Välikangas et al. 2016). For the second analysis, integrating multiple phosphoproteomics datasets from distinct varieties independently generated, we added a batch-effect correction step using ComBat (Johnson et al. 2007), implemented for proteomics using the HarmonizR package from R (Voß et al. 2022) coupled with a prior-imputation step using the Random Forest Decision Tree module MissForest (Stekhoven and Bühlmann 2012; Kokla et al. 2019; Jin et al. 2021) with 100 iterations. Next, to filter out MS-assigned values instead of imputed values, we applied a filtering step to retain phosphosites identified in 75% of the maximum number of replicates per genotype/condition, rounding up. The resulting data frame was used for the statistical analysis.

### Statistical analysis

The dataset consisting of phosphorylated sites was organized into a DataFrame, with each column representing a distinct experimental group or condition. Each row in the DataFrame corresponds to a specific phosphorylated site, with the observed values under different experimental conditions. For Cadenza, the statistical significance of the observed differences in phosphorylation states was assessed using one-way ANOVA for the vsn-transformed intensity values of each phosphorylated site, with a significance threshold set at p ≤ 0.05 using Scipy.stats (Virtanen et al. 2020). For the comparative analysis of phosphoproteomes at 24 °C and 34 °C across different wheat varieties, a two-way ANOVA was performed followed by post-hoc multiple comparisons to determine significant differences (p < 0.05) between genotypes and temperature conditions (**Extended Data table 7**). Additionally, we implemented batch-effect correction and imputation steps to handle missing values and ensure the robustness of our data among distinct LC-MS/MS runs.

### Network inference and kinases/phosphatase database construction

Network inference was performed using NetPhorce in R (Van den Broeck et al. 2023). This package draws upon the principles of dynamic Bayesian networks and includes two stages: 1) pinpointing possible regulator-target pairings, and 2) evaluation and ranking of these pairings based on Bayesian principles. Next, according to the vsn-normalized intensity values and their phosphorylation/dephosphorylation dynamics along the temperature gradient, the percentage fold change between temperatures required to be labelled as an increase/decrease in phosphorylation was set to ≥ 50%, with a minimum fold change threshold set to 50% and *q*-value for the test set to < 0.001. Additionally, to include *a priori* identification of kinases and phosphatases in the network inference, a list of kinases and phosphatases was built using the IWGSC RefSeq v1.0 database for *Triticum aestivum var.* Chinese spring (International Wheat Genome Sequencing Consortium (IWGSC) 2014). The conserved protein kinase domain (PF00069) was used as a guide. To achieve this, we obtained a consensus alignment sequence from the PFAM website, built an HMM profile, and performed an HMMER search using HMMER v3.3.2 (Eddy 1998). We obtained a list of 5914 potential kinases in wheat. Next, we added an additional validation layer for the presence of the classic protein kinase domain using InterProScan 5 (Jones et al. 2014) and confirmed its homology. Unlike kinases, which share a unique conserved protein kinase domain, phosphatases have greater sequence variation. Therefore, orthology-based identification was based on the well-annotated *Oryza sativa* spp. Japonica (International Rice Genome Sequencing Project 2005) and *Arabidopsis thaliana* (Cheng et al. 2017) phosphatases. Known phosphatase identifiers for these species were retrieved from PLAZA (Van Bel et al. 2022), and the best potential ortholog was determined using the four evidence instances from the platform [(1) - tree-based ortholog, (2) orthologous gene family, (3) anchor point, and (4) best-hits-and-inparalogs family]. A total of 475 phosphatases were identified in *T. aestivum*. The generated networks were visualized using Cytoscape 3.8.0 (Shannon et al. 2003).

### UMAP analyses and clustering

We employed Uniform Manifold Approximation and Projection (UMAP) (McInnes et al. 2018), implemented via the umap-learn package in Python, as a critical tool for dimensionality reduction and visualization of complex proteomic data. The number of neighbors and metrics were set to default in all runs, regardless of grouping by samples or previously filtered phosphorylated sites. Additionally, we incorporated soft clustering techniques, specifically Fuzzy C-Means (Bezdek et al. 1984), to identify inherent patterns and groupings within the data with the number of clusters set to 12. During Fuzzy C-Means clustering, each protein in the dataset was assigned a membership score for each cluster, reflecting the degree to which the protein belonged to the cluster.

### Feature selection

For feature selection, we used the random forest regressor function from the Sklearn module in Python (Pedregosa et al. 2012). We allocated 80% of the dataset for training and 20% for testing, setting the random_state to 42 for reproducibility. The model, run with 1000 estimators, was trained to predict the mean phosphosite measurement across all temperatures using phosphosite data as input features. This approach allowed the model to learn the relationship between the input features and the target variable. After training, the model calculated feature importance scores, identifying key phosphosites significant for understanding adaptive responses to different temperatures.

### Data and material availability

The raw mass spectrometry proteomics data, MaxQuant settings, MaxQuant outputs, and sample mapping generated in this study were deposited in the ProteomeXchange Consortium via the PRIDE database under accession code PXD047670 [username: - Password: 4wzGQAdt]. All the python code (Jupyter notebooks) for generating Figs. 1 and 2 as well as the individual subpanels and dataset are available at: https://github.com/Cassio-Lima/Fonseca_de_Lima_2023.

## AUTHOR CONTRIBUTIONS

C.F.F.d.L. and I.D.S. conceived the project. L.V.d.B. designed the algorithms and the databases. C.F.F.d.L., T.Z., B.v.d.C. performed analysis and benchmarks and developed the software. R.S. and A.M.L. provided guidance and input for the analyses. I.D.S. supervised the project. C.F.F.d.L., and I.D.S. wrote the manuscript. All authors reviewed and approved the final manuscript. The authors declare no conflict of interest.

## ACKNOWLEDGEMENTS

This work was supported by the Research Foundation – Flanders (FWO.OPR.2019.0009.01). TZ was recipient of a PhD grant from the Chinese Scholarship Council (201706910095) and a postdoctoral research mandate from the UGent BOF (BOF22/PDO/113), the Foundation for Food and Agriculture Research (FFAR CA18-SS-0000000026), Benson Hill, VIB, BASF, the United Soybean Board (2020-152-0134), and the North Carolina Soybean Producers Association (20-122) to R.S., A.M.L., and I.D.S.

## SUPPLEMENTARY FIGURES

**Extended data figure 1.**
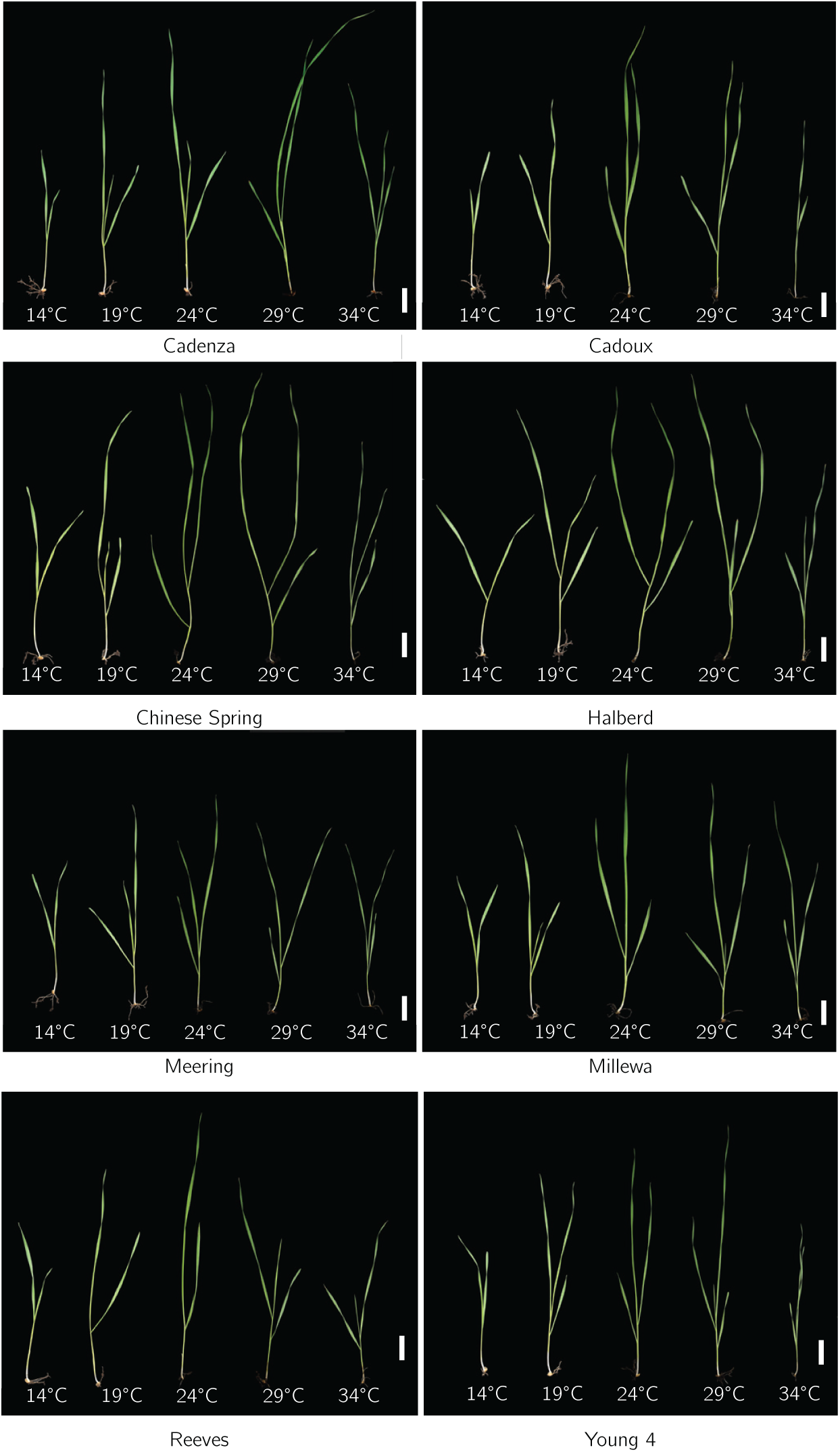
Representative images for the 8 studied wheat varieties along the experimental temperature gradient. White scale bar, 3 cm.

**Extended data figure 2.**
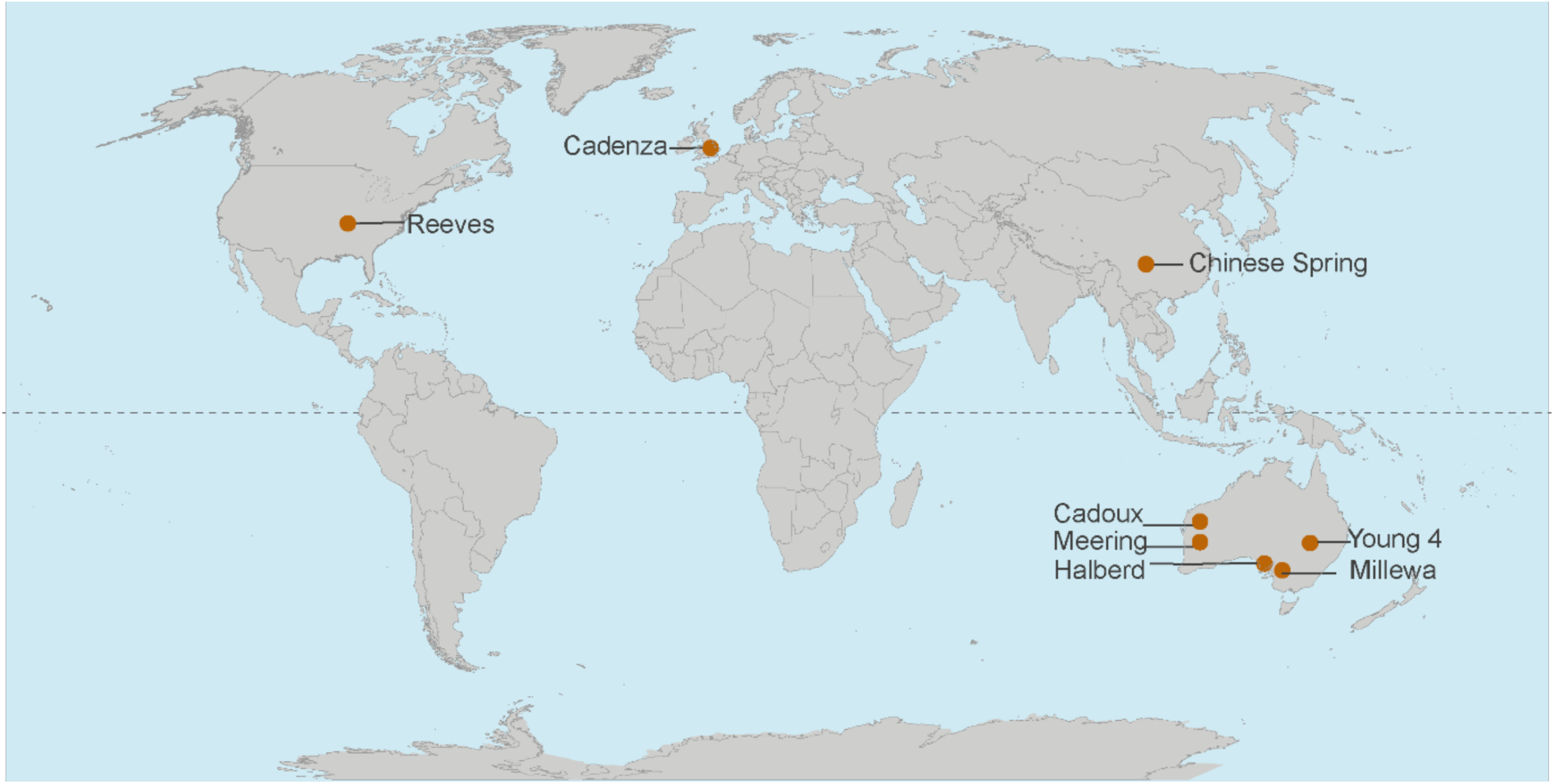
World map showing the provenance of the 8 studied wheat varieties.

**Extended data figure 3.**
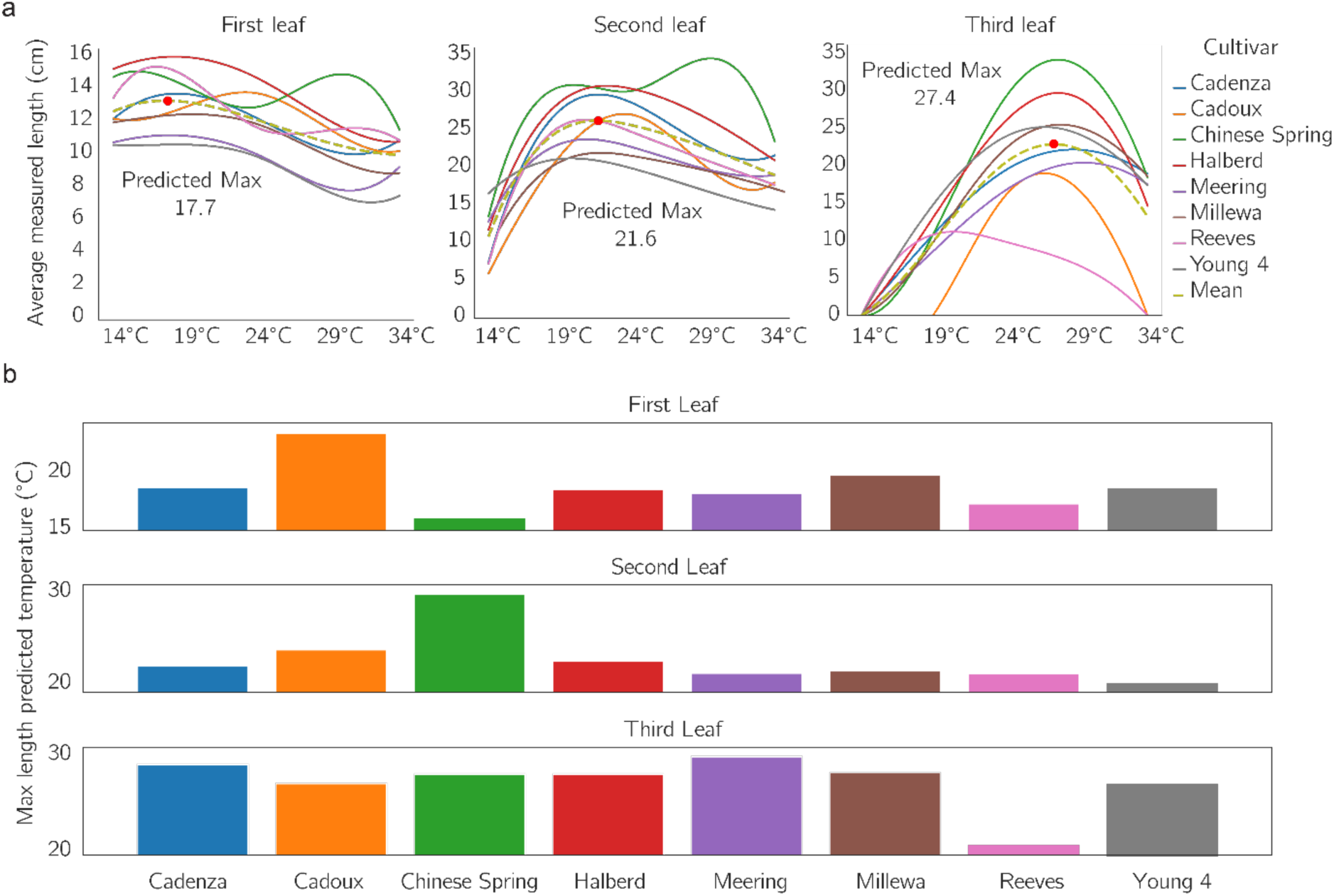
Leaf wheat length along a temperature gradient. a. Third-degree polynomial curve fit from unscaled leaf length data. The dotted line indicates the best fit across all varieties, and red dots indicate the temperature at which leaf length is predicted to reach its maximum. b. Predicted max temperatures per variety. Raw maximum/minimum values are shown.

**Extended data figure 4.**
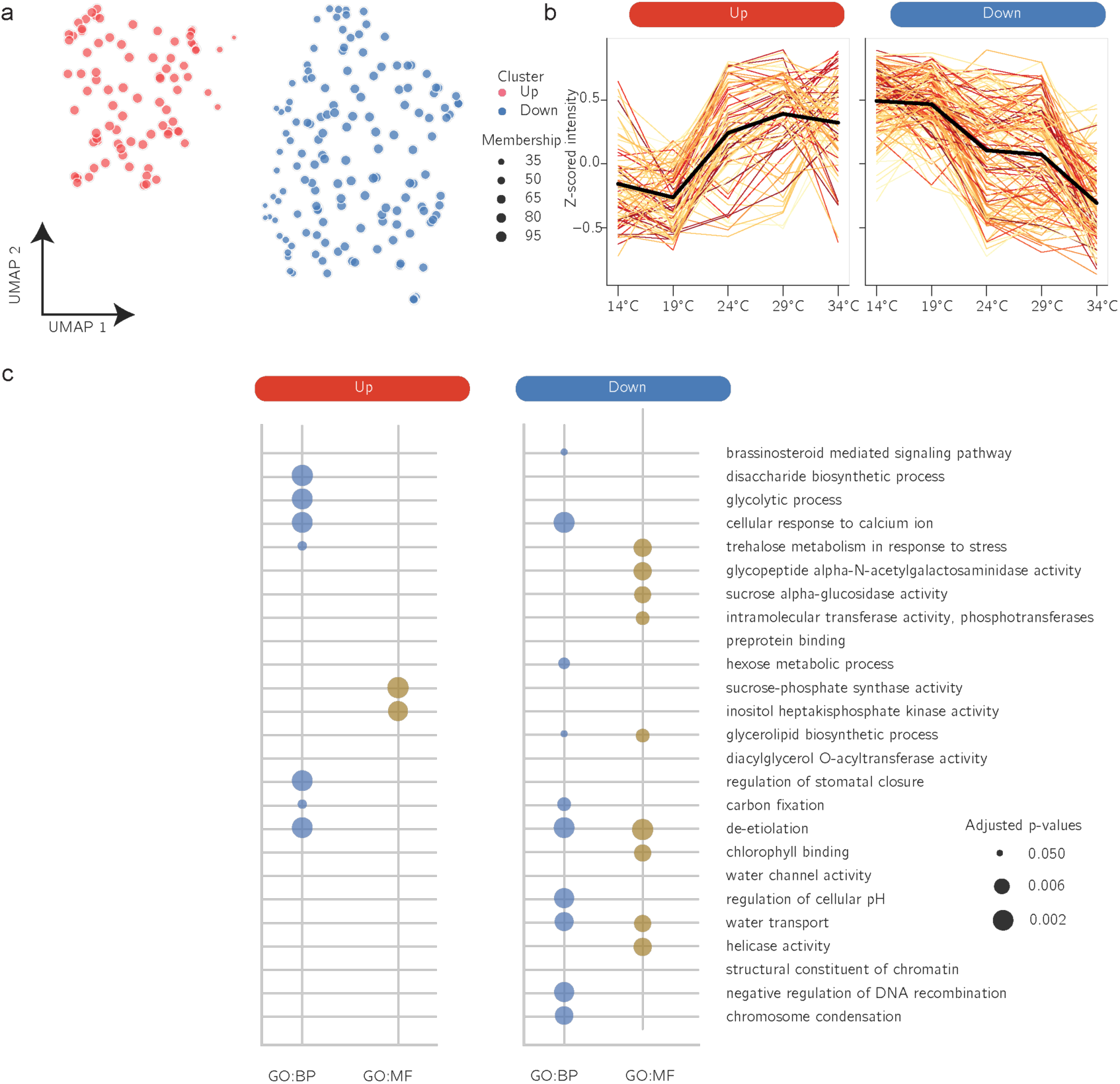
Phosphorylated sites that were regulated in Cadenza (209) along the temperature gradient. a. Soft fuzzy cluster of profiles for the phosphorylated sites revealing a “Up” and “Down” groups according to their trend of phosphorylation along the gradient. Sizes are correlated to their membership to a particular cluster. b. Grouped profiles; each line represents a single phosphorylated sites while the black line best represents the cluster. c. Gene ontology enrichment of the two clusters observed.

**Extended data figure 5.**
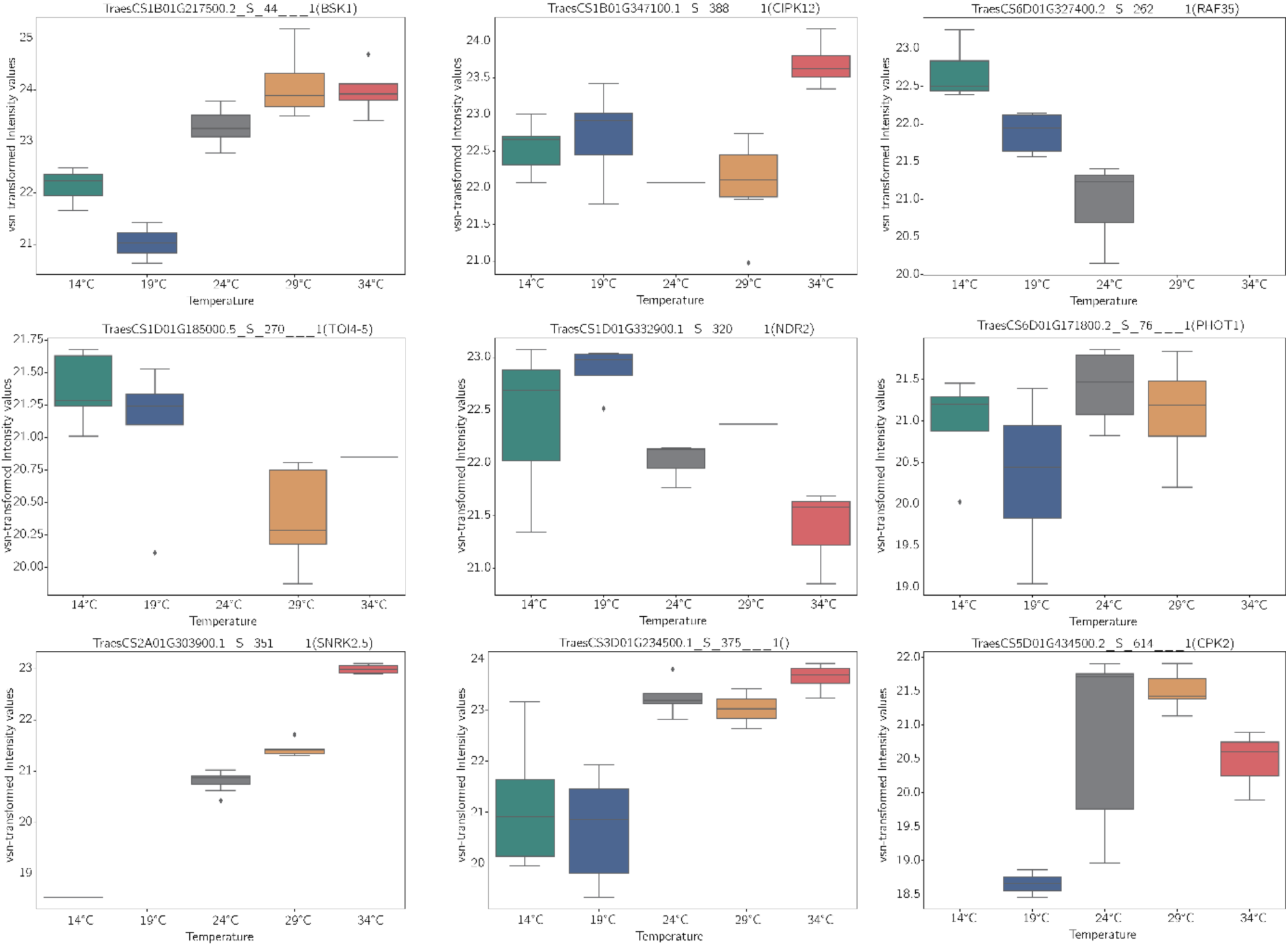
Profile of phosphorylated sites in kinases predicted in the Dynamic Bayesian Network. Wheat identifiers are shown (protein isoform identified by LC-MS/MS, followed by the amino acid residue, position within the protein and multiplicity. Between parenthesis the name of the best-hit Arabidopsis orthologue. Colours represent distinct temperatures. All members of this group have a *p*-value < 0.05 for the ANOVA test.

**Extended data figure 6.**
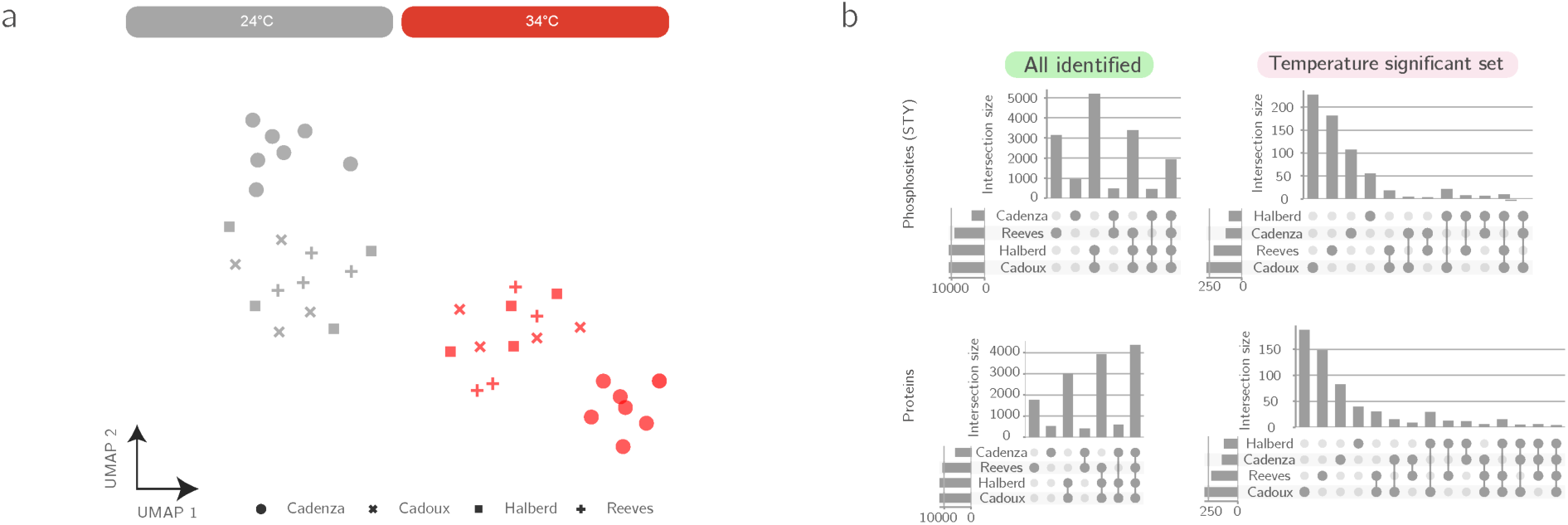
Multiple comparison and data integration of experiments on distinct wheat varieties (Cadenza, Halberd, Cadoux and Reeves). a. Corrected data for 24°C and 34°C second leaf phosphoproteome experiment on the four indicated wheat varieties. b. Comparison of all identified phosphorylated sites and proteins among the multiple experiments and significant sets.

## SUPPLEMENTARY DATA TABLES

**Extended data table 1.** List of *Triticum aestivum* L. (wheat) varieties used in the present study. Including common sowing season, origin, heat-sensitivity (HS) status and literature evidences for classification; pedigree data obtained from the UK wheat varieties pedigree (Fradgley et al. 2019).

**Extended data table 2.** S2A. Measurements (in cm) of distinct leaf numbers (first, second and third) from all the studied varieties after submission to the listed temperature for 14 days. Empty cells represent absence of the organ/no measurement. S2B. Raw intensities for Phospho (STY) Sites in Cadenza pre-filtered/transformed or corrected. S2C. Raw intensities for Phospho (STY) Sites in Cadoux & Halberd pre-filtered/transformed or corrected. S2D. Raw intensities for Phospho (STY) Sites in Reeves pre-filtered/transformed or corrected.

**Extended data table 3.** S2A. Intensity values of 209 high confidence phosphorylated sites that were significantly regulated (One-way ANOVA = *p-value* ≤ 0.05) distributed into different clusters of profiles with respective membership values. Number of cluster was manually set to 12. Index column represented here by “Ptn_AA_pos_multiplicity” represents the identified protein isoform, aminoacid residue, position and multiplicity associated with the quantified value. Values have been ranked from low to high in a scale of red (lowest) to green (highest). S2B. List of significant candidates for indicated pariwise comparision (Student’s T-test, p < 0.05)

**Extended data table 4.** List of kinases with dynamic phosphorylation profile along the temperature treatments with statistical test results for individual varieties.

**Extended data table 5.** S5A. List of phosphorylated sites in Cadenza ranked by feature importance. S5B. List of wheat-Arabidopsis best orthologues.

**Extended data table 6.** Adjusted intensity values for high-temperature (24 °C vs. 34 °C) regulated protein-phosphorylated sites in Cadenza, Cadoux, Halberd, and Reeves.

**Extended data table 7.** Statistical test results (one/two-way ANOVA) for commonly identified phosphorylated sites for Halberd (heat-tolerant) and Cadoux (heat-sensitive).

## Notes

### Competing Interest Statement

The authors have declared no competing interest.

